# Shorter effective lifespan in laboratory populations of *D. melanogaster* might reduce sexual selection

**DOI:** 10.1101/2021.04.09.439133

**Authors:** Avani Mital, Manaswini Sarangi, Bodhisatta Nandy, Neha Pandey, Amitabh Joshi

## Abstract

The role of sexual selection in mediating levels of sexual conflict has been demonstrated in many experimental evolution studies on *Drosophila* spp. where competition among males for mating was the target of selection. Sexual selection has also been shown to affect the evolution of life-histories. However, the influence of divergent life-histories on reproductive strategies and, therefore, sexual selection and possibly sexual conflict, has been less well studied. We examined *D. melanogaster* populations selected for a short development time and early age at reproduction for changes in reproductive behaviour and traits that are proxies of sexual selection. We report a large reduction in reproductive competition experienced by the males of these populations, compared to ancestral populations that are not consciously selected for rapid development or early reproduction, potentially leading to reduced sexual selection. We show that rapidly developing and early reproducing populations have very low levels of mating in their lifetime (females are more or less monandrous), low courtship levels, shorter copulation duration, and longer time from eclosion to first mating, compared to the controls. These results are discussed in the context of the previously demonstrated reduction of inter-locus sexual conflict in these populations. We show that life-history strategies might have a large and significant impact on sexual selection, with each influencing the other and contributing to the complexities of adaptation.

**Significance statement:** Sexual conflict, often manifested as an arms-race between males and females trying to enhance their own reproductive success at some cost to the other, is of great evolutionary interest because it can maintain genetic variation in populations, prevent the independent optimization of male and female traits, and also promote speciation. Sexual selection, or variation in mating success, is well known to affect levels of sexual conflict. However, it is not so clear whether, and how, the regular evolution of life-histories also affects sexual selection. Here, we show that life-history evolution in fruit fly populations selected for traits not directly related to sexual conflict might, nevertheless, mediate the possible evolution of altered sexual conflict levels through effects on sexual selection. Populations that evolved to develop to adulthood fast, and reproduce relatively early in life, are shown to potentially experience less sexual selection, which can explain the low sexual conflict levels earlier observed in them.

## Introduction

Due in part to Darwin’s independent treatment of natural selection (Darwin 1859, 1868) and sexual selection (Darwin 1871), the two sub-fields have largely developed separately, and the causes and consequences of viability/fecundity selection and sexual selection have often been studied by evolutionary biologists with somewhat differing backgrounds and interests. An unfortunate consequence of this outcome has been that studies of life-history evolution and sexual selection did not meaningfully intersect for many decades, even though they both focus on opportunities for, and timing and distribution of, reproductive output, albeit from somewhat different points of view. Parental investment per offspring differs between males and females of most sexually reproducing species (Bateman 1948; Trivers 1972), resulting in different reproductive strategies for male or female fitness maximization, often leading to differences in optimal mating rates for males and females (Bateman 1948; Andersson 1994). Such differences can give rise to inter-locus sexual conflict (Parker 1979; Chapman et al. 2003; Anrqvist and Rowe 2013), resulting in arms-race like dynamics with males evolving to manipulate female reproductive choices, and females, in turn, evolving to circumvent such manipulation (Chapman et al. 2003; Anrqvist and Rowe 2013). Sexual selection and sexual conflict are clearly intertwined and, indeed, have been studied together in some detail over the past few decades (e.g., Wigby and Chapman 2004; Linklater et al. 2007; Edwards et al. 2010; Nandy et al. 2013a, b). However, even though viability and fecundity selection can shape life-histories in ways that can in principle either heighten or reduce sexual selection and, thereby, sexual conflict, life-history evolution has not yet been integrated with studies of sexual selection and sexual conflict in similar detail.

Several studies have provided evidence for sexual selection affecting life histories. For example, Hollis et al. (2017) report the evolution of faster development and maturation in monogamous compared to polygamous *D. melanogaster* populations. In males of the decorated cricket, *Gryllodes pirillas*, increasing reproductive effort with age was shown to correlate with slower ageing and longer lifespan (Archer et al. 2012). Similarly, Zajitschek et al. (2009) demonstrated sexual dimorphism in age dependent patterns of survival and reproduction in the field cricket *Teleogryllus commodus*, likely due to differences is age-dependent reproductive strategies between males and females. Recently, evidence for the evolution of longer development times and decreased desiccation and starvation resistance in polyandrous *D. pseudoobscura* lines compared to monogamous lines has also been reported (Garlovsky et al. 2021).

In contrast, the direct influence of life-history changes on sexual selection and sexual conflict has been investigated relatively rarely, even though these influences can potentially have large effects on both sexual selection and sexual conflict. For example, a life-history providing a relatively short duration of time for reproduction can reduce sexual selection by constraining opportunities to re-mate, driving lower levels of competition among males. The latter, in turn, could drive an evolutionary reduction in inter-locus sexual conflict levels. Thus, effective adult lifespan (the duration of adult life in which reproduction can occur), in particular, might alter the level of sexual selection, all else being equal. This can be particularly important in discrete generation laboratory systems such as *Drosophila*, where continued adult lifespan beyond the point at which eggs are collected to initiate the next generation does not contribute to Darwinian fitness. For example, populations of *D. melanogaster* that experienced longer durations of effective adult life evolved increased male offense and defense ability at late ages and an improved ability to induce typical female post mating responses, compared to their controls (Service 1993; Service and Fales 1993; Service and Vossbrink 1996). Similarly, selection for early reproduction in seed beetles *Callosobruchus maculatus* resulted in the evolution of more frequent early-life mating compared to those selected for late reproduction (Maklakov et al. 2010).

One of the clearest demonstrations of life-history evolution affecting sexual conflict has come from a study showing considerably reduced sexual conflict in populations of *D. melanogaster* subjected to long term selection for rapid development and early reproduction in the laboratory relative to control populations from which they were derived (Ghosh and Joshi 2012; Mital et al. 2021). This was shown to be driven in part by the much smaller body size of flies from the selected populations (Mital et al. 2021), a consistent correlated response to strong selection for rapid development in *D*. *melanogaster* across multiple studies (Zwaan et al. 1995; Nunney 1996; Chippindale et al. 1997a; Prasad et al. 2000). The observation that size reduction in these selected populations likely drove the evolution of reduced levels of sexual conflict appears to be consistent with studies showing smaller males to be less harmful to females than larger males (Partridge et al. 1987a, b; Pitnick 1991; Pitnick and Garcia-Gonzales 2002).

However, in addition to being selected for rapid development, which results in reduced body size, the selected populations used by Ghosh and Joshi (2012) were also selected for relatively early reproduction (around day 3 of adult life) compared to controls (around day 11 of adult life). The selected populations, thus, have an effective adult lifespan that is less than one-third that of the controls. With a 3 day effective adult lifespan, there is limited scope of multiple matings by females, especially if smaller, resource deprived females fail to remate readily. This might result in a further reduction in sexual selection and sexual conflict in these populations, beyond that due to the smaller body size of selected population males (Ghosh and Joshi 2012; Mital et al. 2021). Here, we examine this hitherto unexplored aspect of our faster developing populations by asking whether differences in the breeding ecology of these selected and control populations might also be possibly affecting sexual selection.

We looked at various proxies of the strength of sexual selection – lifetime mating frequency, courtship frequency, female maturation time and copulation duration – in our selected and control populations. Mating frequency has been used as a proxy for potential sexual selection previously (Kuijper and Morrow 2009). Courtship frequency is indicative of male mating effort since *D. melanogaster* males perform extensive and energetically expensive courtship for which they likely bear a fitness cost (Cordts and Partridge 1996; Anholt et al. 2020). Moreover, courtship by males potentially provides an opportunity for females to exercise mate choice (Gavrilets et al. 2001; see Anholt et al. 2020 for review). Copulation duration is another estimate of reproductive effort/investment by males as copulation in *D*. *melanogaster* lasts substantially longer than required for sperm transfer alone, with the additional time spent in the transfer of accessory gland proteins (Acps) (Gilchrist and Partridge 2000). Many of these Acps are known to mediate sexually antagonistic effects in females (Chapman et al. 1995; Wolfner et al. 1999; Chapman 2001). We note that we have not directly assessed sexual selection via variation in male reproductive success and that, therefore, our results are suggestive rather than conclusive.

## Methods

### Study Populations

We used eight large, outbred *D. melanogaster* populations that have a common ancestry. Four of these populations were selected for rapid pre-adult development and early reproduction and are referred to as the **FEJ**s (<u>**F**</u>aster developing, <u>**E**</u>arly reproducing, <u>**J**</u>B derived); the other four were ancestral controls, called the **JB**s (<u>**J**</u>oshi <u>**B**</u>aseline). Details of the ancestry of these populations and their maintenance have been reported previously (JB: Sheeba et al. 1998; FEJ: Prasad et al. 2000).

In summary, JBs are maintained on a 21day discrete generation cycle with eggs collected into 40 replicate vials (9.5 cm ht × 2.4 cm dia) per population (60-80 eggs/ 6mL of banana-jaggery medium). All adults typically emerge by the 12^th^ day from egg collection and are transferred to fresh food vials on days 12, 14 and 16. On day 18, all flies are collected into a Plexiglas cage (25 × 20 × 15 cm^3^) provided with food supplemented with additional yeast for three days, before starting the next generation from eggs laid during an 18 h time window on day 21. The FEJ maintenance is similar except that only the first 20-25% of eclosing adult flies from each vial are collected directly into a Plexiglas cage with food supplemented with additional yeast. After three days, eggs are collected to initiate the next generation. All populations are maintained at a breeding adult number of 1500-1800 flies, under constant light, 25°C ± 1°C and about 90% relative humidity. Thus, only the fastest developing flies in the FEJs make it to the breeding pool and reproduce relatively early, i.e., on day three after eclosion. Consequently, by the time of this study (~600 generations of FEJ selection), FEJs were being maintained on a 10-day discrete generation cycle. Since each of the four FEJ populations has been derived from one JB population, we could account for ancestry by treating FEJs and JBs with matched subscripts as random blocks in the statistical analyses. Standardization (common control-type rearing conditions for a full generation in order to equalize non-genetic parental effects) was done only prior to assaying reproductive maturity and copulation duration in the FEJ and JB populations, as we wanted to assay courtship and mating frequencies under conditions approximating the regular maintenance of the populations. It was not possible to record behaviour data blind as the focal flies from the JB and FEJ populations are easily distinguishable based on their body size.

### Mating and Courtship Frequencies

For this assay, we used flies derived from eggs laid in the running cultures, without standardization, as we wanted to assess breeding ecology differences that FEJ and JB flies experience during their maintenance, including those potentially due to non-genetic parental effects. Since the maintenance regimes and development times of the FEJ and JB populations are quite different, assaying them under conditions closely mimicking their normal maintenance necessitates some differences in the assay protocols for the two sets of populations. For JBs, flies from these assay populations were collected on day 12 from egg-lay, corresponding to the time all flies would have eclosed in their regular cultures, pairing five males and five females per vial with fresh banana-jaggery food medium (12 vials per population) to facilitate courtship observations. Observation vials were again set up on days 14 and 16 as described for day 12. For observing flies in the last two days in cages prior to egg lay, we set up 100 pairs of flies per Plexiglas cage (22 × 18 × 18 cm^3^) with a plate of fresh food medium covered with a paste of live yeast, as in their regular maintenance. We set up three replicate observation cages per population; these were slightly smaller than the maintenance cages, to facilitate behavioural observations. Flies were first collected into a regular cage (from holding vials in case of JBs and culture vials in case of FEJs) and then 100 males and 100 females were lightly anesthetized (using carbon dioxide) and transferred into each replicate observation cage. In case of the fast developing FEJs, only the first ~25% of the eclosing flies (~day 6.5 from eggs) were selected to become part of the observation set and directly shifted to observation cages with a plate of fresh food medium covered with a paste of live yeast, as in their regular maintenance.

We took three ‘instantaneous’ observations every four hours (corresponding to one time point) starting from day one of observation (i.e., day 12 from JB culture initiation (egg-lay), ~day 6.5 from FEJ culture initiation) noting the number of males observed to be performing a courtship behaviour (Spieth 1974) towards a female, and the number of pairs in copula. Observations were conducted over 144 hours (six days) in the vials for only JBs, as the FEJs do not have a corresponding vial stage as adults in their regular maintenance, and for 72 hours (three days) in Plexiglas cages for both JBs and FEJs, for each replicate population. We calculated the daily average of courtship and mating frequency (fraction of males that were courting or mating) for each day. Daily average frequencies were then summed over all days of the effective lifetime, i.e., nine days for JB populations and three days for FEJ populations, each for mating or courtship.

### Copulation Duration and Female Maturation Time

To obtain adults of the same age at the beginning of the assay, FEJ cultures were started about 60 hours after the JB cultures, to account for the difference in their egg to adult development times. We allowed an egg laying time of only 30 minutes, in order to have a narrow eclosion distribution, and collected females that emerged within a one-hour time window. Females were paired with one day old virgin males (single pair per food vial), with 20-25 pairs per population, under each of two conditions (presence or absence of live-yeast supplement on food) and kept undisturbed for continuous observation. In their regular maintenance, the first matings occur in cages in the presence of food supplemented with additional yeast for FEJs, but in vials without yeast for JBs. We recorded time from eclosion till first mating (maturation time) and copulation duration under both yeasted and unyeasted conditions. The duration of observation was for 56 hours, or until 80% of pairs per population, per treatment, had mated, whichever occurred first.

### Statistical Analyses

We used mixed model analyses of variance (ANOVA) for data from all the assays. Average time till maturation for a population, lifetime courtship and mating frequency per population, and proportion of wild-type offspring averaged across females per population were used as the response variables. Tukey’s HSD test was used for comparing individual means and calculating all confidence intervals for post-hoc analyses. Since some data were fractional, we also repeated each analysis for those response variables with arcsine square root transformed values to check if there were any differences obtained in the significance of the fixed factors. Selection regime and yeasted/non-yeasted conditions for time till maturation and copulation duration, and selection regime for courtship and mating rate assays were treated as fixed factors in the analysis. All experiments were performed on four replicate populations of a maintenance regime. Replicate populations with matching numerical subscripts (FEJ_*i*_ and JB_*i*_; *i* = 1-4) were assayed together, and also share the same ancestry. Those pairs were therefore treated as random blocks in the analyses. All analyses were carried out using STATISTICA^™^ using Windows Release 5.0B (StatSoft Inc. 1995).

## Results

### Mating and Courtship Frequencies

The differences between FEJ and JB lifetime mating (*F*=1020.928, *P*< 0.0001) and courtship frequencies (*F*=1372.439, *P*< 0.0001) were statistically significant, with JBs showing many fold higher frequencies of both courtship and mating (Fig. 1a, b; Table 1). When we compared the mating and courtship frequencies only within the cage stage (the last three days of their effective lifespan), courtship frequencies were still significantly greater for JB than for FEJ (*F*=910.352, *P*<0.001), but their mating frequencies did not differ significantly (*F*=4.163, *P*=0.1339; Fig. 1c, d; Table 1). Moreover, although overall courtship frequencies were very low for FEJ, the results from the analysis of data from only within the cages indicated a very low courtship requirement for arousal of FEJ females for copulation, given that mating frequencies in the cage were very similar for JB and FEJ, despite courtship frequencies being much lower in FEJ (Fig. 1c, d). It is also worth noting that both courtship and mating frequencies dropped by almost an order of magnitude for JB when only the cage stage was considered, indicating that most mating in the regular cultures of JBs probably occurs during the vial transfer stages, before the flies from all vials representing one replicate population are collected into a cage.

**Table 1.**
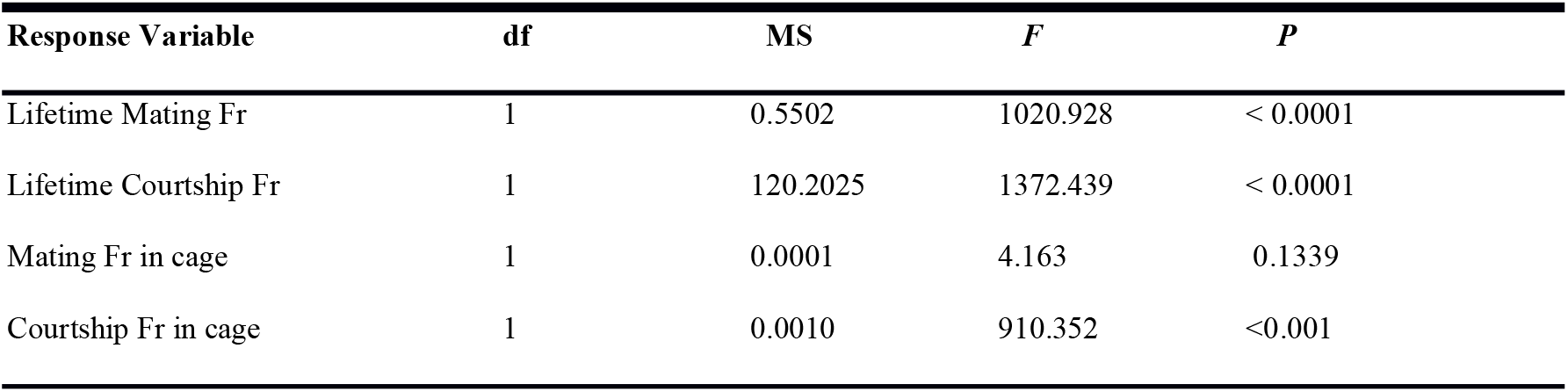
Results summary of ANOVA for effect of selection on mating and courtship frequencies^1^

| Response Variable | df | MS | F | P |
| --- | --- | --- | --- | --- |
| Lifetime Mating Fr | 1 | 0.5502 | 1020.928 | < 0.0001 |
| Lifetime Courtship Fr | 1 | 120.2025 | 1372.439 | < 0.0001 |
| Mating Fr in cage | 1 | 0.0001 | 4.163 | 0.1339 |
| Courtship Fr in cage | 1 | 0.0010 | 910.352 | <0.001 |

**Fig. 1.**
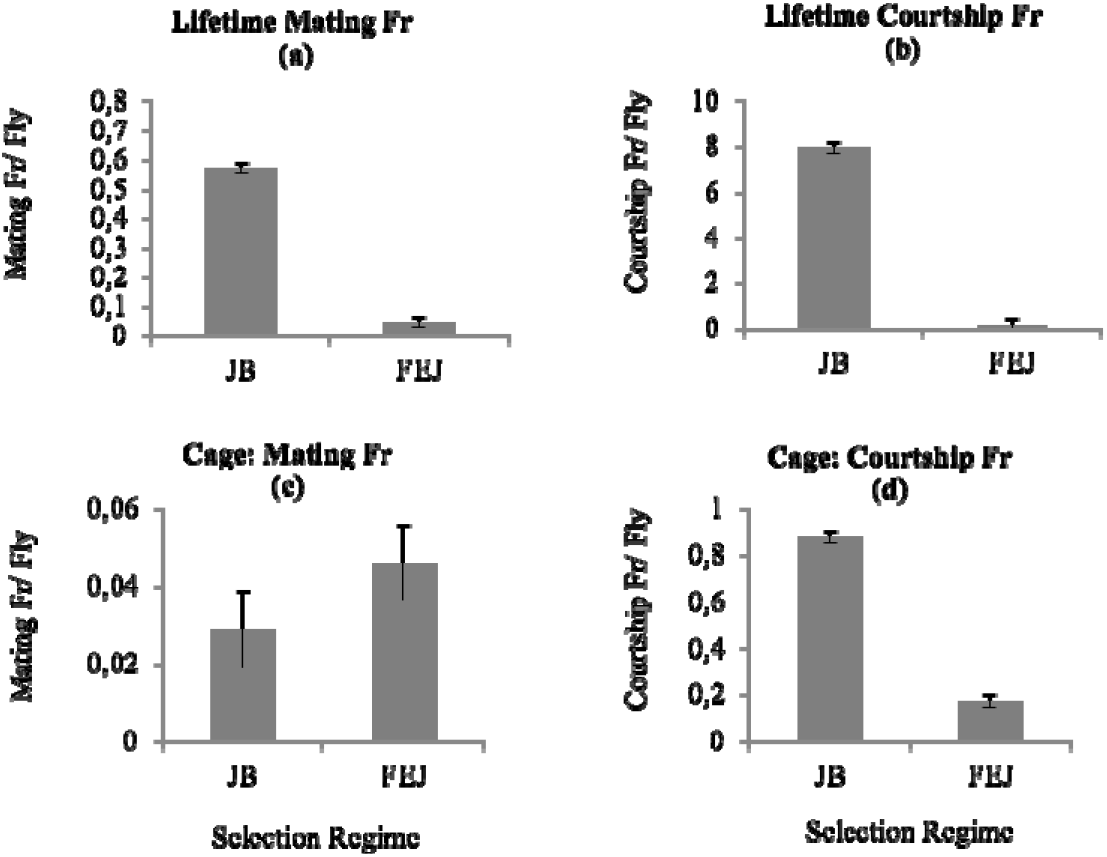
a) Mean lifetime mating frequency averaged across the four replicate populations of JB and FEJ. b) Mean lifetime courtship frequency averaged across four replicate populations of JB and FEJ. c) Mean mating frequency in the cage, during the final three days of effective adult life, averaged across four replicate populations of JB and FEJ. d) Mean courtship frequency in the cage, averaged across the four replicate populations of JB and FEJ. Error bars are 95% confidence intervals around the means.

### Copulation Duration and Female Maturation Time

Maturation time of the JB and FEJ differed significantly (*F*=149.885, *P*=0.0011; Table 2), as did copulation duration (*F*=15.260, *P*=0.0297; Table 3), with JB females taking a shorter time from eclosion to first mating than FEJ females, consistent with previous reports (Prasad 2004; Ghosh-Modak 2009), and mating for longer than FEJ females (Fig. 2a, b). The drastic reduction in FEJ development time, especially pupal duration, may have resulted in the postponement of many aspects of reproductive development to adulthood (Prasad 2004), causing FEJ females to take ~10 hours longer than JB females to mature (Fig. 2a). Although there is considerable evidence for change in mating behaviour with adult diet and nutrition in flies (Fricke et al. 2008; Schultzhaus et al. 2018; Duxbury and Chapman 2020), our results did not demonstrate any effect of additional yeast on female maturation time or copulation duration; there was no significant main effect of environment or of the selection-by-environment interaction (Table 2, 3). Female condition (adults being fed a protein rich diet in this case) is also known to increase male courtship vigour through an increased attractiveness of the female (Long et al. 2009), but such effects on the FEJ or JB males, or differences in these effects, did not manifest in our assays.

**Table 2.**
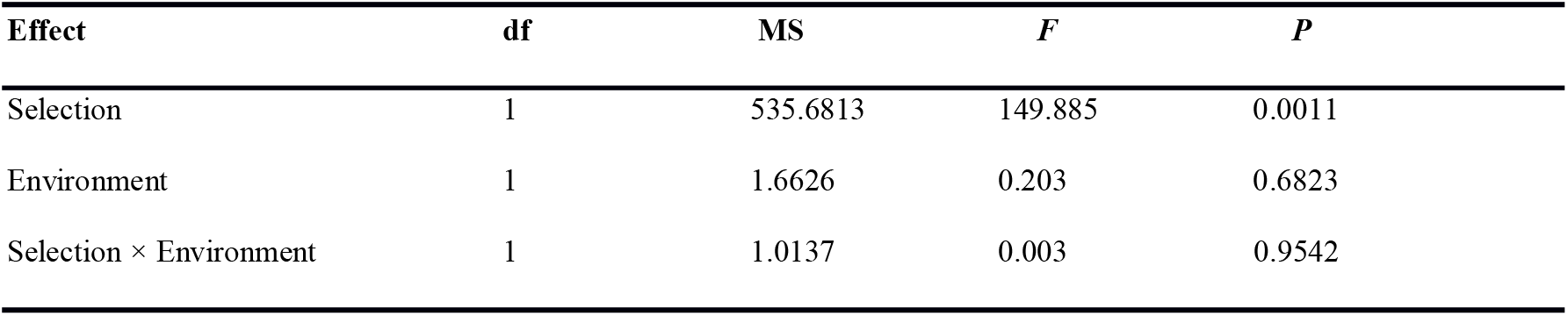
Results summary of ANOVA for maturation time^2^

| **Effect** | **df** | **MS** | ***F*** | ***P*** |
| --- | --- | --- | --- | --- |
| Selection | 1 | 535.6813 | 149.885 | 0.0011 |
| Environment | 1 | 1.6626 | 0.203 | 0.6823 |
| Selection × Environment | 1 | 1.0137 | 0.003 | 0.9542 |

**Table 3.** Results summary of ANOVA for copulation duration^3^

| **Effect** | **df** | **MS** | ***F*** | ***P*** |
| --- | --- | --- | --- | --- |
| Selection | 1 | 143.8742 | 15.260 | 0.0297 |
| Environment | 1 | 0.6389 | 0.6460 | 0.4802 |
| Selection × Environment | 1 | 0.0867 | 0.0326 | 0.8680 |

<sup>3</sup> Selection regime and environment (yeasted or non-yeasted) are fixed factors in the ANOVA. Random factors and interactions cannot be tested for significance in this design, and have been left out for brevity.

**Fig. 2.**
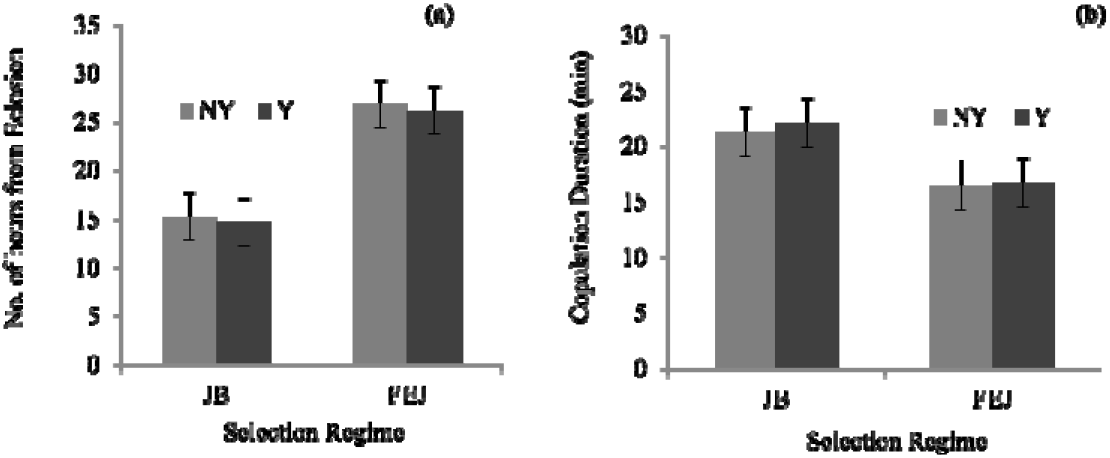
a) Mean maturation time, or mating latency of virgins from eclosion, averaged across four replicate populations of JB and FEJ, under yeasted (Y) and non-yeasted (NY) conditions. b) Mean copulation duration of the first mating averaged across four replicate populations of JB and FEJ, under yeasted (Y) and non-yeasted (NY) conditions. Error bars are 95% confidence intervals around the means.

## Discussion

Sexual conflict is thought to be an incidental by-product of sexual selection (Trivers 1972, Parker 1979) and sexual conflict can be affected by the intensity and scope of sexual selection. Several studies on insects have investigated how reproductive strategies may evolve upon changing their breeding ecology. These studies establish a strong relationship between sexual selection, altered by manipulating the degree of competition among males for mating, and sexual conflict related traits. For example, populations of *Drosophila* sp. were subjected to either monogamy (Holland and Rice 1999; Pitnick et al. 2001; Crudgington et al. 2005; Hollis et al. 2014; Wensing et al. 2017), or different operational sex ratios (Wigby and Chapman 2004; Linklater et al. 2007; Edwards et al. 2010; Nandy et al. 2013a, b) in order to experimentally alter the degree of polygamy and, therefore, sexual selection experienced by the flies. Traits that promote male-specific fitness reduced in the monogamy adapted populations (Holland and Rice 1999; Pitnick et al. 2001; Crudgington et al. 2005; Wensing et al. 2017), and females from these populations experienced greater fitness loss when paired with males adapted to polygamy than to monogamy (Holland and Rice 1999; Crudgington et al. 2005). Similarly, females from populations with male-biased sex ratio had higher resistance to mate harm compared to females from female biased sex ratio populations (Wigby and Chapman 2004; Nandy et al. 2014). In contrast to these studies, sexual selection and sexual conflict in our FEJ populations have been altered as a correlated response to selection on life-history traits like development time and adult age at effective reproduction.

Alterations in life history are also predicted to affect sexual selection and, consequently, the evolution of sexually antagonistic traits (Bonduriansky et al. 2008; Adler and Bonduriansky 2014), and at least a subset of these effects is expected to be through alterations in the breeding ecology of a population. Breeding ecology, which includes not only the timing of reproductive activity, but also various other aspects of reproduction and mating behaviour, such as male-female encounter rate, promiscuity, sexual dimorphism etc., is expected to be defined by the life history strategy adopted by the population. For example, if males in a population have a “live-fast, die-young” life history strategy, male-male competition is expected to be intense, all else being equal (Bonduriansky et al. 2008, and references therein). A few investigations have shown the evolution of sexually selected traits as a consequence of selection for life history traits such as lifespan (Service 1993; Service and Fales 1993; Service and Vossbrink 1996; Makklakov et al. 2010). These reports generally support the idea that evolution of life history traits can result in changes in male reproductive investment, which may further result in reduction in sexual conflict in the population. Yun et al. (2017) showed that the sexual conflict in a population living in a physically complex habitat can be less than that in a simple habitat. If this effect of habitat type is due to females’ access to a spatial refuge, then male-female encounter rate is directly implicated as an important determinant of the level of conflict in that population. Therefore, factors such as, locomotor activity, opportunity of interaction etc. are also expected to be important modulators of the conflict.

Our results complement and extend these studies, with our observations on breeding ecology of laboratory populations of *D. melanogaster* that have earlier been shown to have evolved reduced levels of inter-locus sexual conflict as a correlated response to selection for rapid egg-to-adult development and early reproduction (Ghosh and Joshi 2012; Mital et al. 2021), suggesting a reduced opportunity for sexual selection in these populations. A reduction in the time available for mating in the FEJ populations appears to considerably reduce reproductive competition by constraining re-mating opportunities. Thus, the lower levels of sexual conflict in these populations, compared to their ancestral controls (Ghosh and Joshi 2012), earlier shown to be partly due to their smaller body size (Mital et al. 2021), are also likely to be partly a consequence of reduced sexual selection and male-male competition for mates. Moreover, we also find males from our selected populations to have substantially lower courtship frequency and shorter copulation duration, indicating potentially reduced investment in pre- and post-copulatory competition. Taken together, our present study and those of Ghosh and Joshi (2012) and Mital et al. (2021), to our knowledge, constitute one among very few examples potentially connecting life-history evolution, sexual selection and sexual conflict.

In conditions closely approximating the regular maintenance of the FEJ and JB populations, we found that the FEJ flies have a much lower mating rate than the JB controls. To better interpret the large difference in life-time mating rates between the two sets of populations, we scaled the mating frequency results by the maximum possible number of matings achievable in that time for that population. We multiplied the FEJ mating rate by 24 (maximal possible mating events being once every hour) to obtain an expected number of life-time mating events, which got rounded off to one mating during their effective adult life. In comparison, the projected life-time mating events for a JB fly is six matings during their effective adult life (scaling factor of 12, mating once every two hours). These scaling factors were chosen based on the mating duration and likely recovery required by FEJ and JB males before attempting re-mating (A Mital, personal observation) and we believe these are overestimates of the maximum number of matings possible within a day. Therefore, one and six are conservative estimates of the expected lifetime mating events per fly for FEJs and JBs, respectively.

One limitation of our assay is that it only tells us that FEJ adults in their regular cultures are, on an average, probably getting to mate just once, as compared to an average of six matings for JB adults in their cultures. In the absence of data on the variance of male mating success in these populations, it could be speculated that the FEJ populations are likely monandrous, since females are typically refractive after a mating and tend not to re-mate for quite some time, whereas some males could be obtaining multiple matings and others none at all. In that case, although post-copulatory sexual selection would be low, there might be substantial pre-copulatory sexual selection on males in the FEJ populations. However, we believe this is unlikely to be the case in the FEJ populations, for the following reason.

There has, to our knowledge, been only one study of the variance of male mating success in large, outbred *Drosophila* populations under assay conditions approximating their running cultures (Joshi et al. 1999). That study was carried out on the B populations of Rose (1984), which are the ultimate ancestors of our JB control populations but differ in having been maintained on a 14 day discrete generation cycle with no cage maintenance at any life-stage. Joshi et al. (1999) examined the mating success of individual marked males over the relevant part of adult life. Except for early eclosing males that found themselves in vials with a female-biased sex-ratio, there was no evidence for a non-Poisson distribution of male mating success (Joshi et al. 1999). Even those early eclosing males showed a non-Poisson distribution of male mating success in only three out of five replicate populations, and in each of those populations, the variance in male mating success was less than half of the mean. Early eclosing males, thus, were obtaining a large number of matings, due to the availability of more than one female per male, but mating success varied even less than expected through chance alone if all males had equal probabilities of mating (Joshi et al. 1999).

If we examine the life-history in the FEJ, JB, B and UU (first described in Shiotsugu et al. 1997) populations, we can predict that it is likely that the FEJ populations have both reduced pre- and post-copulatory sexual selection, compared to the JB controls. The B populations (Rose 1984) are ancestral to the rest; the UU were derived from the B in 1991 and shifted from a two week to a three week discrete generation cycle (Shiotsugu et al. 1997), the JB were derived from the UU in 1997 and remained on a three week cycle (Sheeba et al. 1998), and the FEJ were derived from the JB in 1998 and subjected to selection for rapid development and early reproduction (Prasad et al. 2000). The B populations were not specifically selected for rapid development by truncating the eclosion distribution. However, compared to the UU populations, the B populations showed reduced development time (Shiotsugu et al. 1997); the development times of the UU and JB populations are comparable, though they were never assayed together. The development time data coincide well with the effective adult life span of these various sets of populations: FEJ are the fastest developing and have an effective adult life of about 3.5 days, followed by the B with an effective adult life of about 5 days, and then the UU and JB that are the slowest developing and have an effective adult life of about 11 days. In the B populations, the number of matings obtained by a male in its effective lifetime ranged from 0 to 7, with a mean of 1.8 (data in Table 1, Joshi et al. 1999). This mean mating success, based on direct observations of marked males, is considerably lower than what we infer for the JB controls from our assay (6 lifetime matings), but higher than what we infer for the FEJ (1 lifetime mating). Given these observations on the B populations, which have somewhat rapid development and short effective lifespan, we expect that is likely that males in our FEJ populations are undergoing considerably less pre- and post-copulatory sexual selection as compared to the JB controls. However, an unequivocal conclusion of reduced sexual selection in the FEJ populations would require a direct assessment of variation in male reproductive success in the FEJ and JB populations.

Another result indicating considerable change in reproductive behaviour of our selected populations is the large reduction in their lifetime courtship frequency, compared to controls (Fig. 1b), which persisted even in the cage stage (last three days of effective adult life, Fig. 1d) whereas similar differences in mating frequency were not seen in the cage stage (Fig. 1c). There are multiple plausible causes for this result, as we discuss below.

The period between eclosion and first reproduction (maturation time) is typically the only opportunity for adult females to feed and thereby accumulate additional resources for egg production (Luckinbill et al 1985; Chippindale et al. 1997b). Such compensatory feeding would be especially important for FEJ females given their small size and low lipid content at eclosion (Prasad 2004). The already resource limited FEJ females are expected to then reduce all energetically expensive activities, which may render increased receptivity selectively advantageous. That FEJ males are significantly less harmful and females less resistant to mate harm has already been reported earlier (Ghosh and Joshi 2012, Mital et al. 2021). Moreover, frugal courtship by males and a low arousal threshold in females is likely to be selectively advantageous also because mating at least once is necessary to have any fitness, and the time available for reproduction in the FEJ breeding ecology is short and further reduced by a long maturation time, (Fig. 2a). In addition, since FEJs have also evolved a significantly different developmental schedule and maturation time (Prasad 2004; Ghosh-Modak 2009; Fig. 2a), developmental changes may further contribute to poor courtship by males, especially since males complete reproductive activity at a very young adult age of about three days. For instance, in some studies on *D. melanogaster*, males have been found to keep maturing reproductively, especially in terms of accessory gland growth, for up to six days after eclosion, with younger males tending to have poor sperm competitive ability and courtship effort as compared to more mature males (Ruhmann et al. 2016). Finally, males from the controls JB populations may be selected for high courtship effort during the final three days of their adult life, as they have about 80% paternity share of the offspring that form the next generation if they mate with females during the ‘cage’ stage (Supplementary Material, Fig. S1) (Fig. 2c, d). JB males would gain a high fitness reward for mating effort in this time despite low chance of mating success. This may also indicate relatively high variation in male mating success in the control population, a long-standing argument for stronger selection in males for exaggerated reproductive traits (Bateman 1948; Andersson 1994).

More generally, however, low courtship frequencies despite similar mating frequencies among males that we see in the cage stage (Fig. 1c, d) can also be explained in the context of the chase-away selection model for the evolution of exaggerated sexual traits in *D. melanogaster* (Holland and Rice 1998; Gavrilets et al. 2001). The model suggests that if persistent male mating effort is harmful to females (sexual conflict) then high female resistance towards such sensory manipulation will evolve. This may be interpreted as increased female bias/preference, in turn selecting for higher courtship effort from males (Holland and Rice 1998, 1999). Since persistent male courtship is known to reduce female survival in *D. melanogaster* (Partridge and Fowler 1990), reduced sexual conflict driven by low male-male competition in FEJs may release the sexes from this co-evolutionary cycle, and appear as lack of female choice and male courtship effort. Therefore, low courtship frequency in FEJs, nevertheless resulting in a mating frequency similar to that of the JBs in the last three days of effective adult life, may be driven by female ecological constraints, changes in development, reduced male-male competition, or a combination of these. Put together, our results on the lifetime mating and courtship frequencies in the FEJ populations suggest the possible evolution of reduced sexual selection and conflict as a consequence of strong selection for rapid development and an extremely short effective adult life.

The interplay between life-history and sexual selection is expected to ultimately determine the optimum investment strategy for a population. In our study populations, the changes that have possibly resulted in reduced sexual selection would also be expected to release the flies from certain reproductive investment demands. We observed that FEJ flies mate for around 15 minutes on average, compared to about 25 minutes for JB flies (Fig. 2b). Production of Acps is known to be costly to males (Chapman and Edwards 2011) and it has been suggested that in *D. melanogaster*, most sperm transfer occurs during the first half of mating (Gilchrist and Partridge 2000), the remaining time being spent in the transfer of Acps responsible for the female post-mating response. We speculate that, reduced energy investments in courtship and in the non-sperm components of the ejaculate in the FEJs would not only explain their shorter copulation duration (Fig. 2b) but potentially also permit the evolution of even smaller flies, further pushing the boundaries of rapid development for which they experience strong selection. Changes in resource use in FEJs have been demonstrated previously; FEJs have evolved larger egg size (B M Prakash and A Joshi, unpublished data) and more eggs produced per unit dry weight by females (Prasad 2004; Ghosh-Modak 2009) than JBs. It is therefore likely that the optimum investment strategy for male FEJs has diverged from that of the ancestral controls. This is further supported by the observations that greater starvation resistance per unit lipid has evolved in the FEJs than JBs (Prasad 2004), and that Acp genes in young FEJ males are downregulated compared to the JBs (Satish 2010; K M Satish, P Dey and A Joshi, unpublished data).

## Conclusion

Our results, taken together with those from the studies of Joshi et al. (1999), Ghosh and Joshi (2012), and Mital et al. (2021), strongly suggest that the changed breeding ecology of our fast developing and early reproducing FEJ populations might have reduced both pre- and post-copulatory sexual selection and, consequently, facilitated the earlier observed evolution of lower inter-locus sexual conflict experienced by them, as compared to their controls. The shorter effective adult life of FEJs, together with strong directional selection for rapid development, has resulted in small flies that appear to exhibit substantially reduced competition among males for mating. These observations complement other studies that have reported reduced sexual conflict upon directly selecting for different levels of male-male competition (Introduction) and exemplify the potential impact of selection on life-history related traits (development time and age at reproduction) on levels of sexual selection and conflict via major changes in reproductive behaviour. This is especially relevant as differences in sexual selection and sexual conflict are often implicated in the emergence of speciation phenotypes (Parker and Partridge 1998). We note that the FEJs and JBs have diverged in their reproductive behaviour to a degree that incipient reproductive isolation has occurred (Ghosh and Joshi 2012). Our work, therefore, also highlights that selection on traits not directly associated with sexual conflict can, nonetheless, drive the evolution of reproductively isolating mechanisms, either directly through rapid development (Mital et al. 2021) or indirectly by affecting sexual selection, or both. Dissecting out such nuanced interactions of life-history and reproductive strategy in affecting sexual selection and sexual conflict is important for understanding the myriad evolutionary consequences that may accompany adaptation to specific ecological challenges.

## Acknowledgment

We are grateful to Sajith V.S, Rajanna N., and Muniraju P. for help with the experiments, and two anonymous reviewers for helpful feedback. This work was funded by Jawaharlal Nehru Centre for Advanced Scientific Research (in-house funding to AJ), the Science and Engineering Research Board (SERB) JC Bose fellowship (recipient: AJ), and in part by personal funds of AJ. JNCASR provided doctoral fellowships to AM, MS, and NP, and a postdoctoral fellowship to BN.

## Footnotes

1 Summary of results of a one-way ANOVA each for lifetime and cage mating and courtship frequencies. Main effects of selection regime for the two traits are shown. In this design, random factors and interaction effects cannot be tested for significance and have been left out for brevity.

2 Selection regime and environment (yeasted or non-yeasted) are fixed factors in the ANOVA. Random factors and interactions cannot be tested for significance in this design, and have been left out for brevity.

3 Selection regime and environment (yeasted or non-yeasted) are fixed factors in the ANOVA. Random factors and interactions cannot be tested for significance in this design, and have been left out for brevity.

