## Supplementary Material for "Shorter effective lifespan in laboratory populations of *D. melanogaster* might reduce sexual selection" (from another version of this preprint: version 3, 2022-01-02)

Supplementary Material 1

Breeding Success of JB Males

Earlier studies have shown that the timing of mating can be a source of variation in reproductive success among *D. melanogaster* females (Long et al. 2010), implicating age-specific mating and courtship patterns as crucial factors for understanding the breeding ecology of populations with a discrete generation cycle. Assessment of overall mating and courtship rates alone, without information on fertilization success, would grossly ignore the details of male-female post-copulatory interactions that can affect final fitness outcomes. The breeding ecology of the ancestral control populations (i.e., JBs) has a relatively prolonged effective adult life (~11 days) and fitness is attained through production of progeny by the females at the end of this period in an 18-hour oviposition window. Since flies are reproductively active from a very early age (as early as 12 hours from eclosion, Fig. 2a), it is reasonable to expect matings to be distributed across the entire effective adult lifespan, perhaps non-uniformly. In addition, *D. melanogaster* shows strong last male sperm precedence (Lefevre and Jonsson 1962) and complex interactions between sperm and the female anatomy that influence paternity success (Miller and Pitnick 2002) . Therefore, it is important to find out the contribution that mating at different ages has on fertilization success of the eggs laid during the relevant oviposition window. It is important to note here that this issue is greatly simplified in the populations selected for early reproduction (FEJs) where effective breeding life is reduced to about three days. Consequently, we also attempted to quantify age-specific fertilization success in the control population males, to assess which period of their adult life yielded the maximum fertilization of eggs laid on the day that the next generation was initiated.

**Methods**

For examining the breeding success of JB males, a marked strain was needed, and for this we used **SE**, an outbred population like the JBs fixed for the autosomal recessive marker - **s**carlet **e**ye (st). The SE population was created by introgressing the scarlet eye mutation into a mixed JB population maintained on a 21-day discrete generation cycle but on corn-meal molasses medium. For this study, we reared the SE population on banana-jaggery medium for two generations (common control-type rearing conditions for a full generation in order to equalize non-genetic parental effects) prior to assaying. We used females from the SE population, which, upon mating with a JB male (wild type eye colour) will reproduce wild-type offspring. Thus, by housing SE females with SE males overall, but allowing males from the wild type JB population to mate with them during certain phases of their adult life, we were able to assess the age-specific reproductive success of the JB males.

We collected flies from both SE and JB cultures for the experiment at the same time and on day 12 from egg-lay and set up 10 males and 10 females in each breeding vial. We divided the effective adult lifespan (from eclosion to egg-collection) of the JB flies into four phases based on the vial transfer and cage collection regime as experienced by JBs in their regular maintenance protocol. For Phase 1 (P1), we introduced JB males with SE females on day 12 and housed them together till day 14, after which the JB males were replaced with SE males of the same age using light carbon dioxide anesthesia. Similarly, we housed SE females with JB males only between days 14 and 15 (P2), 16 and 17 (P3) and 18, 19 and 20 (P4). At all other times, SE females were housed with SE males. After day 20, the SE females were allowed to lay eggs for 18 hours. We set up 32 oviposition vials for each phase P1-P4, sampling five females from each of the breeding vials. The SE females were allowed to lay eggs (one female per vial) for 18 hours before being discarded. The vials were maintained at 25ºC ± 1ºC and constant light for the next 10-11 days until all flies had eclosed. We quantified the proportion of wild-type progeny for each female (breeding success of the JB males) and averaged this across females for a particular phase. We estimated breeding success of males from all four replicate JB populations.

### Results

A significant effect of phase (df=3, MS=0.53, $F$=121.625, $P$<0.001) indicated overwhelming last male precedence in reproductive success in the JB populations, showing that mating in the final three days of their effective lifespan resulted in as much as 80% paternity share of eggs laid on day 21 from egg collection (Fig. S1). The first two days of mating (P1) yielded the least breeding success, significantly lower than that of the two middle phases (corresponding to day 14 to day 18 from egg collection), with there being no significant difference between phases two and three, as revealed by pair wise comparisons using Tukey's HSD (Fig. S1).

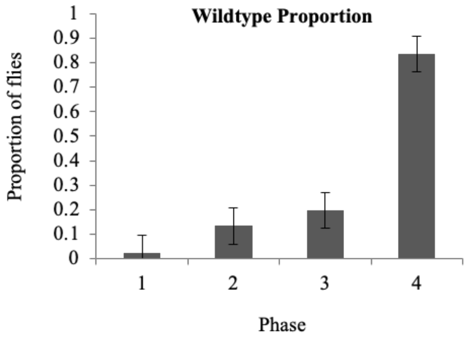


Fig. S1 Mean proportion of wild-type flies among the progeny of the SE flies resulting from eggs laid on day 21, from each of the four phases in the breeding success assay, averaged across the four replicate JB populations. Error bars are 95% confidence intervals around the means. The results show that the JB males have the greatest fertilization success when they mate during Phase 4, immediately before egg-collection on day 21.

### Discussion

The results from the breeding success experiment provide further evidence that high mating effort (courtship frequency) is under positive selection in the JB populations, especially in the cage stage during the last three days of effective adult life, despite mating frequency being very low (Fig. 1c, d). JB populations show significant last male fertilization success, with mating in the last three days resulting in ~80% paternity share (Fig. S1) of eggs collected to initiate the next generation, indicating a high fitness reward for mating effort in this time despite low chance of mating success.. The greater paternity share from matings occurring in the last window can also be explained by females having already laid eggs that were fertilized by earlier matings, but in a discrete generation population those earlier laid eggs do not contribute to fitness. Clearly, substantial sexual selection is experienced by JB flies as males experience very intense competition, especially in the final few days of their effective adult life, unlike FEJ males.
